# TMEM16F activation by Ca^2+^ triggers plasmalemma expansion and directs PD-1 trafficking

**DOI:** 10.1101/249227

**Authors:** Christopher Bricogne, Michael Fine, Pedro M. Pereira, Julia Sung, Maha Tijani, Ricardo Henriques, Mary K Collins, Donald W. Hilgemann

**Affiliations:** UCL Cancer Institute, University College London, Gower St, London, UK.; University of Texas Southwestern Medical Center, Department of Physiology, Dallas, Texas, USA; MRC Laboratory for Molecular Cell Biology, University College London, Gower St, London, UK; National Institute for Biological Standards and Control, Blanche Lane, South Mimms, Herts, UK; Present Address since September 2016, Okinawa Institute of Science and Technology, Onna-son, Okinawa, Japan

## Abstract

TMEM16F, an ion channel gated by high cytoplasmic Ca^2+^, is required for cell surface phosphatidylserine exposure during platelet aggregation and T cell activation. Here we demonstrate in Jurkat T cells and HEK293 cells that TMEM16F activation triggers large-scale surface membrane expansion in parallel with lipid scrambling. Following TMEM16F mediated scrambling and surface expansion, cells undergo extensive membrane shedding. The membrane compartment that expands the cell surface does not involve endoplasmic reticulum or acidified lysosomes. Surprisingly, T cells lacking TMEM16F expression not only fail to expand surface membrane, but instead rapidly internalize membrane via massive endocytosis (MEND). The T cell co-receptor PD-1 is selectively shed when TMEM16F triggers membrane expansion, while it is selectively internalized in the absence of TMEM16F. Its participation in this trafficking is determined by its single transmembrane domain. Thus, we establish a fundamental role for TMEM16F as a regulator of Ca^2+^-activated membrane trafficking.

## Introduction

Eukaryotic cells sequester phosphatidylserine (PS) into the cytoplasmic leaflet of the plasma membrane (PM) (Leventis and Grinstein, 2010). PS becomes exposed in the outer PM leaflet during extended cytoplasmic Ca^2+^ signaling and the execution of apoptosis (Bevers and Williamson, 2010).

Recent work indicates that PS exposure in response to Ca^2+^ elevation is mediated by the Ca^2+^-activated ion channel, TMEM16F (or Anoctamin 6) (Suzuki et al., 2013; Yu et al., 2015), a widely expressed member of the TMEM16 family (Schreiber et al., 2010). In platelets, this type of PS exposure is an important step in blood coagulation, and the overt phenotype caused by TMEM16F mutations in humans is a bleeding disorder known as Scott Syndrome (Suzuki et al., 2010; Yang et al., 2012). In contrast, PS exposure during apoptosis, which marks cells for removal by phagocytosis (Marino and Kroemer, 2013), is not dependent on TMEM16F and may not require Ca^2+^ signals (Suzuki et al., 2010).

Crystal structures of a fungal TMEM16F orthologue reveal a hydrophilic groove (Brunner et al., 2014) that might serve as a conduit for both ions and lipids to pass through the TMEM16F channel, and reconstitution experiments using the fungal TMEM16F support this interpretation (Malvezzi et al., 2013; Brunner et al., 2014). In agreement with this model, a 35 amino acid “scrambling” domain identified in TMEM16F is part of this hydrophilic cleft (Gyobu et al., 2015; Yu et al., 2015). These findings are closely consistent with TMEM16F acting as both an ion channel and lipid “scramblase”, whereby both ions and lipids permeate the channel pore.

In addition to its role in PS exposure, effects of TMEM16F activity on membrane trafficking have been reported. TMEM16F is involved in microvesicular ectosome release from platelets and neutrophils (Headland et al., 2015), and in macrophages TMEM16F is necessary for phagocytosis stimulated by the ATP receptor P2×7 (Ousingsawat et al., 2015a). Mice lacking TMEM16F also show deficits in bone development (Ousingsawat et al., 2015b), a process that has been linked to microvesicles (Anderson, 2003). Microglia lacking TMEM16F demonstrate defects in process formation and phagocytosis (Batti et al., 2016). Finally, mice lacking TMEM16F suffer from immune exhaustion, at least in part due to a failure to down-regulate PD-1, leading to a failure to clear viral infections (Hu et al., 2016).

We have described previously that PS exposure can occur in close association with large exocytic responses activated by elevating cytoplasmic free Ca^2+^ to greater than 10 μM in fibroblasts(Yaradanakul et al., 2008). We therefore examined the role of TMEM16F in Ca^2+^-induced membrane expansion. We describe here in both Jurkat T cells and HEK293 cells how cytoplasmic Ca^2+^ elevations cause rapid PM expansion in close parallel with PS exposure and the opening of TMEM16F channels. The absence of TMEM16F blocks both exocytosis and PS exposure. Surprisingly, the absence of TMEM16F promotes massive endocytosis (MEND) (Lariccia et al., 2011) over the same time course in which membrane expansion would have occurred.

Given the novel observation that TMEM16F controls large-scale membrane trafficking events, we next examined whether specific PM proteins traffic with the corresponding membrane movements. We show in T cells that the immune response regulator PD-1 is selectively shed in vesicles after TMEM16F activation, while it is selectively internalized in the absence of TMEM16F during MEND. This preferential trafficking is dependent on the transmembrane region of PD-1, indicating that the intramembrane segment of PD-1 promotes its participation in membrane domains that can be either shed as ectosomes or internalized via MEND. These results provide a potential explanation for the TMEM16F regulation of PD-1 cell surface expression changes described previously (Hu et al., 2016). As PD-1 is a negative regulator of the immune system, which can be hijacked by various cancers to evade intrinsic immune responses, antibodies that block its function are considered potent cancer therapeutics (Pardoll, 2012). Thus, the Ca^2+^-and TMEM16F-dependent regulation of PD-1 expression in the PM may be of immunological and clinical significance.

## Results

### TMEM16F regulates large-scale Ca^2+^-activated PM expansion

To delete TMEM16F in Jurkat T cells we employed a CRISPR/Cas9 lentiviral vector. We confirmed by genomic DNA sequencing that all TMEM16F copies in Jurkat T cells were mutated, leading to proteins truncated in Exon 2 or 5 (**Figure 1–figure supplement 1**). Commercial TMEM16F antibodies were not reliable for Western blotting, whether employing whole lysates or membrane preparations. Therefore, we employed proteomic analysis of membrane preparations to confirm the quantitative loss of TMEM16F in Jurkat cells (**Figure 1–figure supplement 2**). To determine whether TMEM16F is required for Ca^2+^ mediated PS exposure, we treated cells with the Ca^2+^ ionophore ionomycin and performed fluorescence-activated cell sorting (FACS). Cells with increased cytoplasmic Ca^2+^ were detected via an increase of Fluo4 fluorescence, and cells with PS exposure were dectected via Annexin V staining (**Figure 1**). WT cells responded to ionomycin by correlated increases of cytoplasmic Ca^2+^ and Annexin V binding. Cytoplasmic Ca^2+^ increased in TMEM16F-null cells to the same extent as in WT cells, but TMEM16F-null cells failed to externalize PS, confirming the role of TMEM16F in Ca^2+^-activated PS scrambling (Hu et al., 2016). The Exon 2 truncated TMEM16F-null line was employed in subsequent experiments.

**Figure 1.**
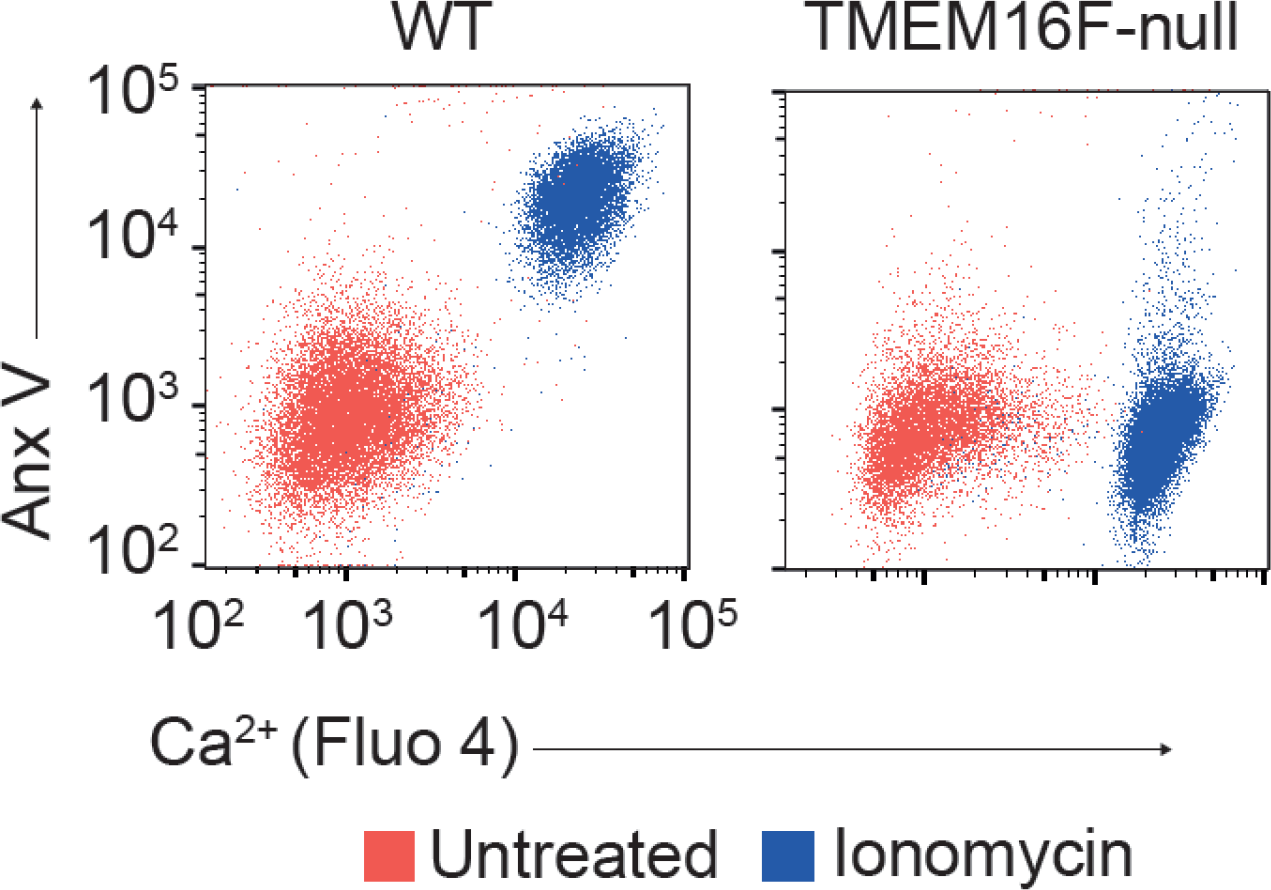
Fluo4-AM loaded WT and TMEM16F-null Jurkat T cells were treated with 5μM Ionomycin for 15 min, then stained with Annexin V and counted using fluorescence-activated cell sorting (FACS).

For real-time analysis of the PM in Jurkat cells, we employed simultaneously patch clamp recording and confocal imaging. PM conductance (G_m_) and capacitance (C_m_) were monitored via continuous square wave voltage perturbation. Membrane area is proportional to C_m_, while G_m_ quantifies the voltage-dependent permeation of ions through the membrane. The surface binding of an impermeable fluorescent membrane dye, FM4-64, was monitored optically. During a 20 second exposure of Jurkat T cells to the Ca^2+^ ionophore, ionomycin (5μM), the PM area doubles on average (**Figure 2A**, C_m_ red trace), and FM4-64 dye binding increases by nearly three-fold (**Figure 2A**, **Figure 2-figure supplement 1**). FM4-64 binding reverses rapidly when the dye is removed, indicating that the dye does not significantly enter cells (**Figure 2A-4**). In contrast to canonical exocytosis, which indeed occurs in T lymphocytes (Matti et al., 2013), membrane expansion in these experiments was insensitive to VAMP2 cleavage by tetanus toxin and did not require cytoplasmic ATP or (**Figure 2C**). Simultaneous with the onset of membrane expansion, the PM conductance of Jurkat T cells increased substantially and declined nearly to baseline in parallel with the asymptotic rise of C_m_ (**Figure 2A**. G_m_, blue trace). Relationships between ion conduction via TMEM16F and membrane expansion are analysed in an accompanying article (Fine, XXXX).

**Figure 2.**
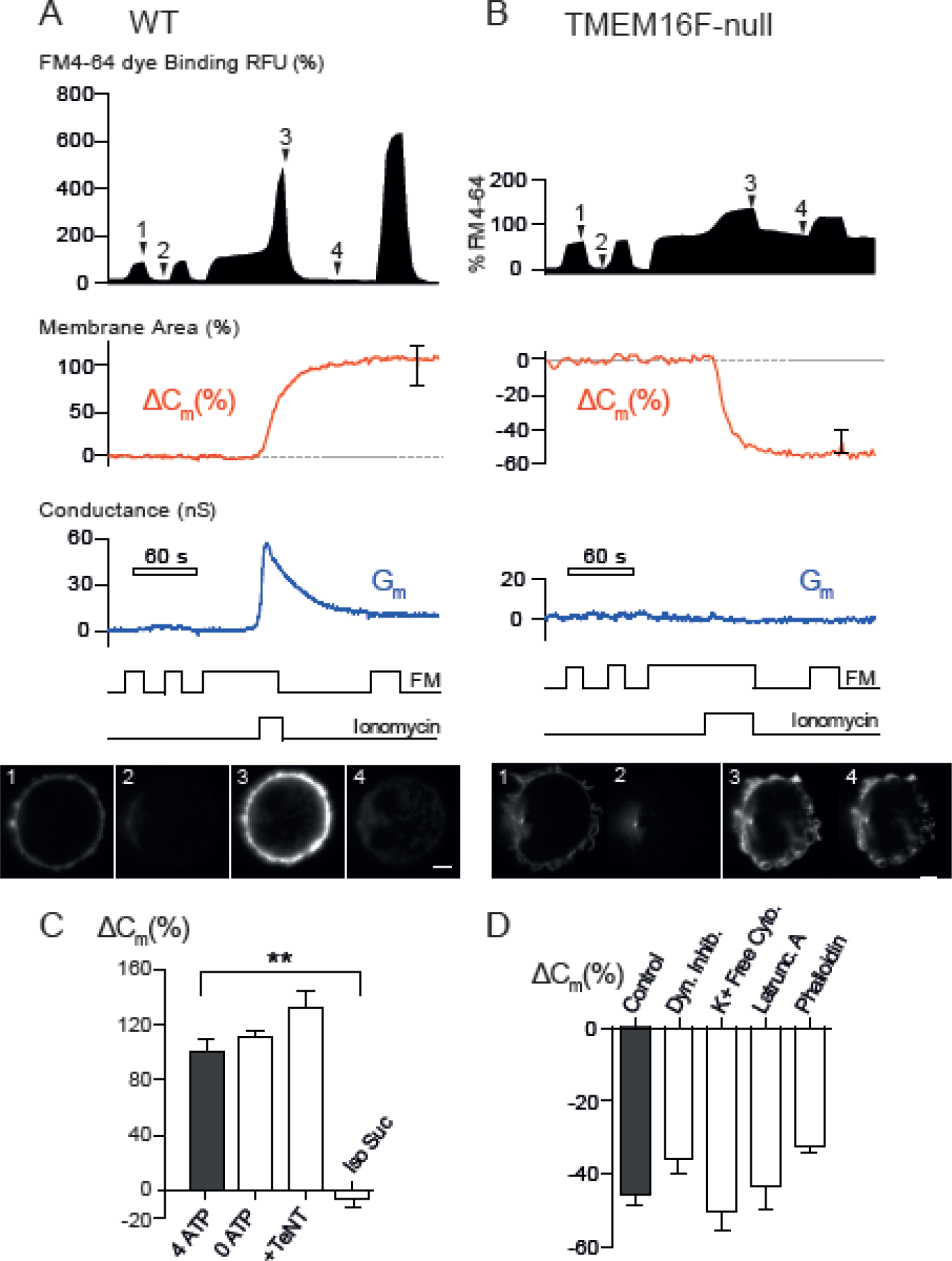
**(A)** Whole-cell patch clamp recordings indicating real-time measurements of membrane area (Cm) and transmembrane ion mobility (conductance, Gm) in WT Jurkat T cells with concurrent confocal imaging using standard Ringer and cytoplasmic solutions. FM4-64 (5μM) and ionomycin (5μM) were applied and washed as indicated. Single experimental traces with SEM values at 150 sec. post-stimulus display membrane area and FM4-64 fluorescence as a percentage of baseline values with confocal images at indicated time points. n=10. Scale bar is 5 um. **(B)** The same protocol as in (A) for TMEM16F-null Jurkat cells. n=10. **(C)** Percentage change in C_m_ 30 s after ionomycin treatment of WT Jurkat cells. Cells were perfused with standard extracellular and cytoplasmic solutions plus indicated conditions for more than 200 s before ionomycin treatment. Neither 400 nM Tetanus Toxin L chain (TeNT) nor ATP/GTP exclusion (0 ATP) showed any significant effect on ionomycin stimulated C_m_ changes. (**p < 0.01) Isotonic replacement of most ions with sucrose blocked membrane expansion **(D)** Percentage change in C_m_ 30 s after ionomycin treatment of TMEM16F-null Jurkat: Results are from left to right: KCl/NaCl solutions (Control), 3 μM cytoplasmic dynamin inhibitory peptide, symmetrical K+-free NmAs solutions, symmetrical 3 μM Latrunculin A, and 10 μM cytoplasmic Phalloidin. (C, D represent mean values with SEM, n>7)

In TMEM16F-null cells, the Ca^2+^-activated increase of conductance was abolished (**Figure 2B**, blue trace). Likewise, instead of expanding by two-fold the PM area decreased by 50% over 25 seconds (**Figure 2B**, red trace). The fact that this decrease reflects extensive endocytosis is established unambiguously by the fact that FM4-64 binding does not reverse after dye is washed off. Rather, the dye becomes locked into membrane structures below the cell surface (**Figure 2B-4, Figure 2–figure supplement 2**). These observations are all consistent with the occurrence of a rapid form of Ca^2+^ activated MEND that we have described previously in fibroblasts (Lariccia et al., 2011). Similar to MEND, these endocytic responses are not blocked by inhibiting clathrin or dynamin, nor by perturbing the actin cytoskeleton with latrunculin or phalloidin (**Figure 2D**).

Further evidence that TMEM16F activity initiates membrane expansion is provided in **Figure 3**. Our previous work demonstrated that cytoplasmic polyamines, such as spermidine and spermine, can switch Ca^2+^-induced membrane expansion responses to MEND responses in fibroblasts (Lariccia et al., 2011), very similar to the effects of TMEM16F deletion. As shown in **Figure 3A**, spermine at physiological concentrations (200 to 500 μM) also switches the Jurkat WT Ca^2+^ response to MEND. Impressively, the TMEM16F conductance is blocked by spermine in close parallel with the switch to MEND, and we point out that spermine has been described to block phospholipid scrambling in red blood cells that lack internal membranes (Bratton, 1994). Accordingly, these data are consistent with the idea that phospholipid scrambling by TMEM16F is linked to the membrane expansion responses.

**Figure 3.**
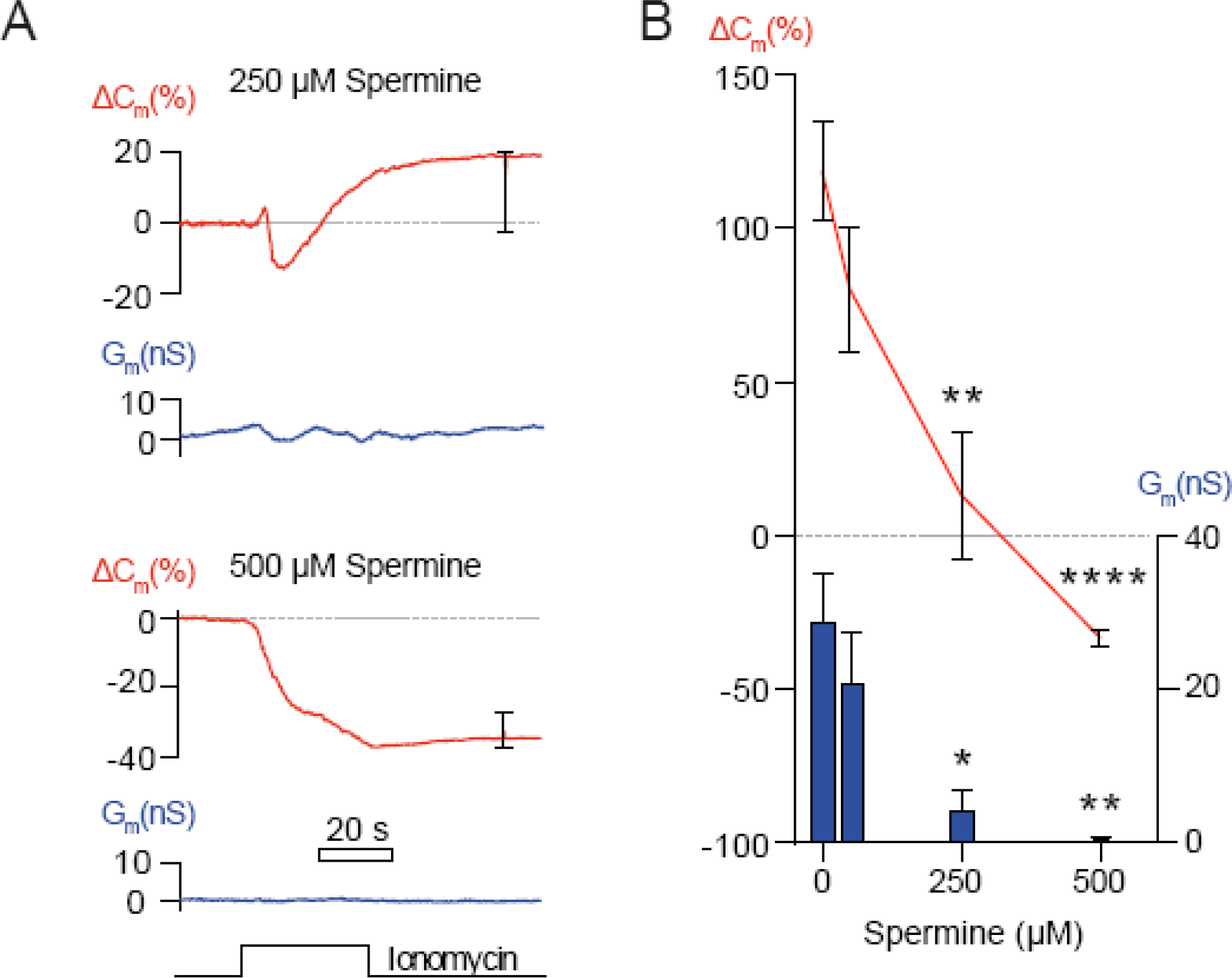
The polyamine, spermine, blocks membrane expansion and TMEM16F conductance with similar concentration dependencies. **(A)** Spermine at 250 and 500 μM was added to cytoplasmic solutions. Cells were held for at least 300s before ionomycin treatment. Single experimental traces are given with mean C_m_ change and SEM. n>7. **(B)** Change in membrane area (ΔC_m_ in %) and peak G_m_ magnitudes were determined during ionomycin application. C_m_ and G_m_ responses were compared to control responses without spermine (^*^ p ≤ 0.05, ^**^ p ≤ 0.01, ^***^p ≤ 0.001 ^****^p ≤ 0.0001).

### The origins of membrane that expands the cell surface

We next set out to determine what intracellular compartments could contribute to such a large expansion of the PM. We looked for evidence of fusion of recycled intracellular vesicles in Jurkat T cells by utilizing super-resolution confocal microscopy projections of intact cells and PM surface imaging with Total Internal Reflection Fluorescence microscopy (TIRF). Jurkat cells were incubated with FM4-64 for one hour to label internalized endosomes, washed to remove extracellular dye, and attached to fibronectin coated dishes. **Figure 4A** describes 0.75 μm z-plane sections projected onto a 2D micrograph to reveal significant FM dye labelling of sub-micron vesicles throughout the cytoplasm. After five minutes of treatment with ionomycin there was no significant change in the number of particles observed indicating that the majority of internally labelled recycled membrane does not participate in our observed membrane expansion (**Figure 4B**). Interestingly, we routinely observed rapid movements of punctae directly after ionomycin application. Viewed alongside the lack of a significant change in the total number of punctae and the slow temporal resolution of 3D high resolution microscopy this might indicate that sorting of particles occurs rapidly and a subset of labelled membrane near the surface of the cell may still be involved. These fast-moving labelled punctae may fuse back to the surface, but are masked by the large signal that remains entirely cytoplasmic.

**Figure 4.**
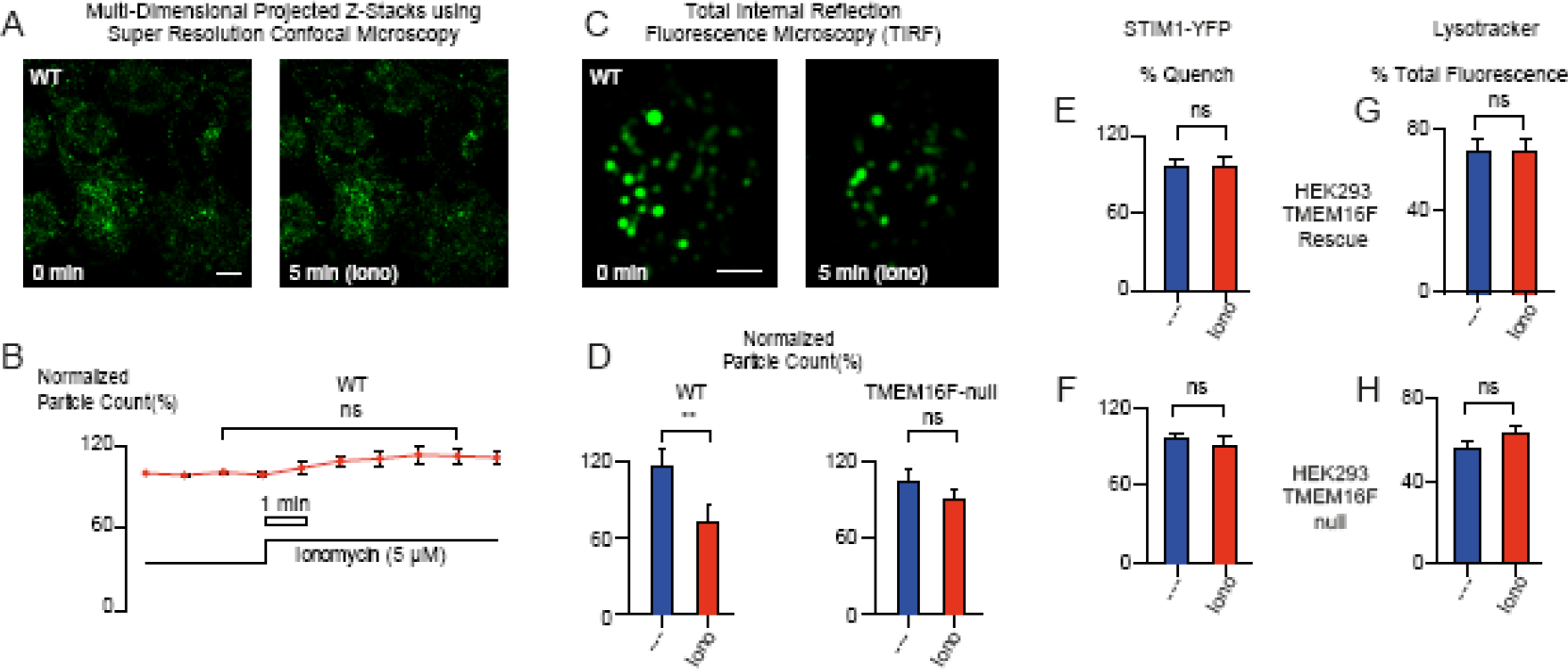
**(A-B)** Multi-Dimensional Super Resolution Airyscan Microscopy. **(A)** Representative pictographs illustrating FM-labelled punctae in WT Jurkat cells before and 5 min after ionomycin treatment. Jurkat cells were attached to fibronectin-coated imaging plates and standard ringer was supplemented with ionomycin after imaging was stabilized for a minimum of 4 frames. 30s frame times **(B)** Quantification of projected 0.75 m z-stacks reveals no significant change to the number of punctae before and after treatment. n=5 independent experiments. **(C-D)** TIRF microscopy captures the Ca^2+^ activated disappearance of FM-labelled puncta in WT but not TMEM16F-null Jurkat T cells following ionomycin treatment. As in **(A-B)** however cells were supplemented with ionomycin 5s before imaging. Values are expressed as a percentage of the initial average number of puncta calculated at 0, 5 and 10 s. **(E-F)** Attached HEK293 TMEM16F-null and mTMEM16F-rescue cells transiently expressing the ER membrane protein STIM1 containing a luminal (ER-resident) YFP fluorophore (Liou et al., 2005) were imaged with extracellular application of 100 g/ml TB to transiently quench extracellular fluorescence of YFP. Cells were exposed to 5μM ionomycin to stimulate membrane fusion. Exposure of the luminal YFP domain of STIM1 during membrane expansion would allow for increased quenching of fluorescence by extracellular TB. No significant change in fluorescence was detected. Analysis shows percentage of initial YFP fluorescence prior to and 300 s after ionomycin stimulation. n=5. Mean values and SEM depicted. **(G-H)** Newly expanded membrane is not associated with lysosomal probes. TMEM16F expressing and TMEM16F-null cells were stained for 60min with 100 nM of LysoTracker^™^ Red. The probe is highly selective for acidic organelles and accumulates in lysosomal compartments. Cells were imaged before and after buffer was supplemented with 5μM ionomycin and 2mM Ca^2+^. Lysosomal fusion to the PM would result in release of fluorescent cargo and loss of labelling. Analysis reveals no detectible loss of fluorescent punctae across both cell lines using ImageJ (NIH) to determine average fluorescence value per cell area with SEM before and after ionomycin stimulation. n=10.

To examine if a subset of vesicles near the surface could account for the large amounts of novel membrane exposed during TMEM16F activation, we labelled Jurkat and Jurkat TMEM16F-null cells with FM 4-64, as before, and analysed punctae loss using TIRF microscopy, as previously described (**Figure 4C)** (Zhang et al., 2007). For WT cells, the number of FM-dye stained punctae per cell significantly decreased after Ca^2+^-ionophore by 30% over five minutes. DMSO treated control cells displayed a 14% increase in the number of punctae observed. (**Figure 4D**). TMEM16F-null cells showed no significant loss of punctae in response to Ca^2+^-ionophore treatment versus DMSO (**Figure 4D**). While results are consistent with TMEM16F in some way promoting the fusion of recycling endosomes, the extent and rate of events visualized via TIRF fall far short of accounting for the doubling of membrane area detected as an increase in Cm (**Figure 2A**). Thus, additional membrane must contribute to the large PM expansion observed.

Lysosomal compartments are described to fuse to the cell surface during cell wounding (Reddy et al., 2001), and the endoplasmic reticulum (ER) represents a very large repository of intracellular membrane close to the cell surface. To determine if our observed increases in cell surface involved fusion of these intracellular compartments to the PM we expressed fluorescently tagged STIM1 (an ER resident protein) and used LysoTracker to label acidified lysosomes. If ER membrane became contiguous with the surface, the luminal YFP tagged to STIM1 would become exposed to the extracellular solution. Once exposed, the presence of the impermeable quencher Trypan Blue in the extracellular buffer would reduce YFP fluorescence intensity (Wan et al., 1993). Likewise, Lysotracker^®^ is a soluble fluorescent compound that accumulates in acidified compartments. Upon lysosomal-PM fusion, the lysosomal pH would rapidly increase in addition to outward diffusion of the dye causing a loss of Lysotracker fluorescence intensity. **Figure 4E-H** shows that neither ER nor lysosomal membranes participate significantly in TMEM16F mediated exocytosis. In this connection, we have described previously similarly large Ca^2+^ activated membrane expansion in BHK fibroblasts. Ultrastructural analysis indicated that vesicular structures could not account for the membrane expansion, and the magnitudes of individual ‘fusion’ events were often equivalent to hundreds of secretory vesicles (Wang and Hilgemann, 2008). Taken together, previous reports of large ‘fusion’ events and failure to implicate recycling endosomes, ER, and acidified lysosomes suggest the involvement of membrane networks or other larger membrane structures in membrane expansion, as outlined in an accompanying article (Fine, XXXX).

### TMEM16F exocytosis and scrambling are followed by vesicle shedding

**Figures 5A** and **5B** describe the relationships between cytoplasmic Ca^2+,^ PS exposure, and membrane expansion during ionomycin exposure. **Figure 5A** illustrates concurrent imaging of cytoplasmic Ca^2+^ via the indicator, Fluo-4 (green), and PS exposure (red) via binding of a rhodamine labelled cationic peptide, heptalysine (K7r). We employed K7r, rather than AnnexinV, because it binds anionic phospholipids much faster and does not require Ca^2+^ for binding (Yaradanakul et al., 2008). Exposure to ionomycin (5 μM) results in very large increases of cytoplasmic Fluo-4 fluorescence followed by cell surface labelling of K7r, indicating large increases of cytoplasmic Ca^2+^ lead to externalization and exposure of PS. The full video of this experiment can be seen on **Figure 5-figure supplement 2**. **Figure 5B** illustrates parallel measurements of membrane expansion and phospholipid scrambling, monitored via C_m_ (red trace) and K7r (green trace), respectively. The two signals can be closely overlaid with one another during membrane expansion. Subsequent to membrane expansion and PS exposure, the two fluorescent signals decline in parallel. The fact that the K7r signal is not maintained on cells, as Cm declines, is indicative of the occurrence of extensive membrane shedding. This was readily visible in confocal microscopy and was typically preceded by the development of distinct membrane protrusions **(Figure 2-figure supplement 1**, **Figure 5-figure supplement 1 & 2**)

**Figure 5.**
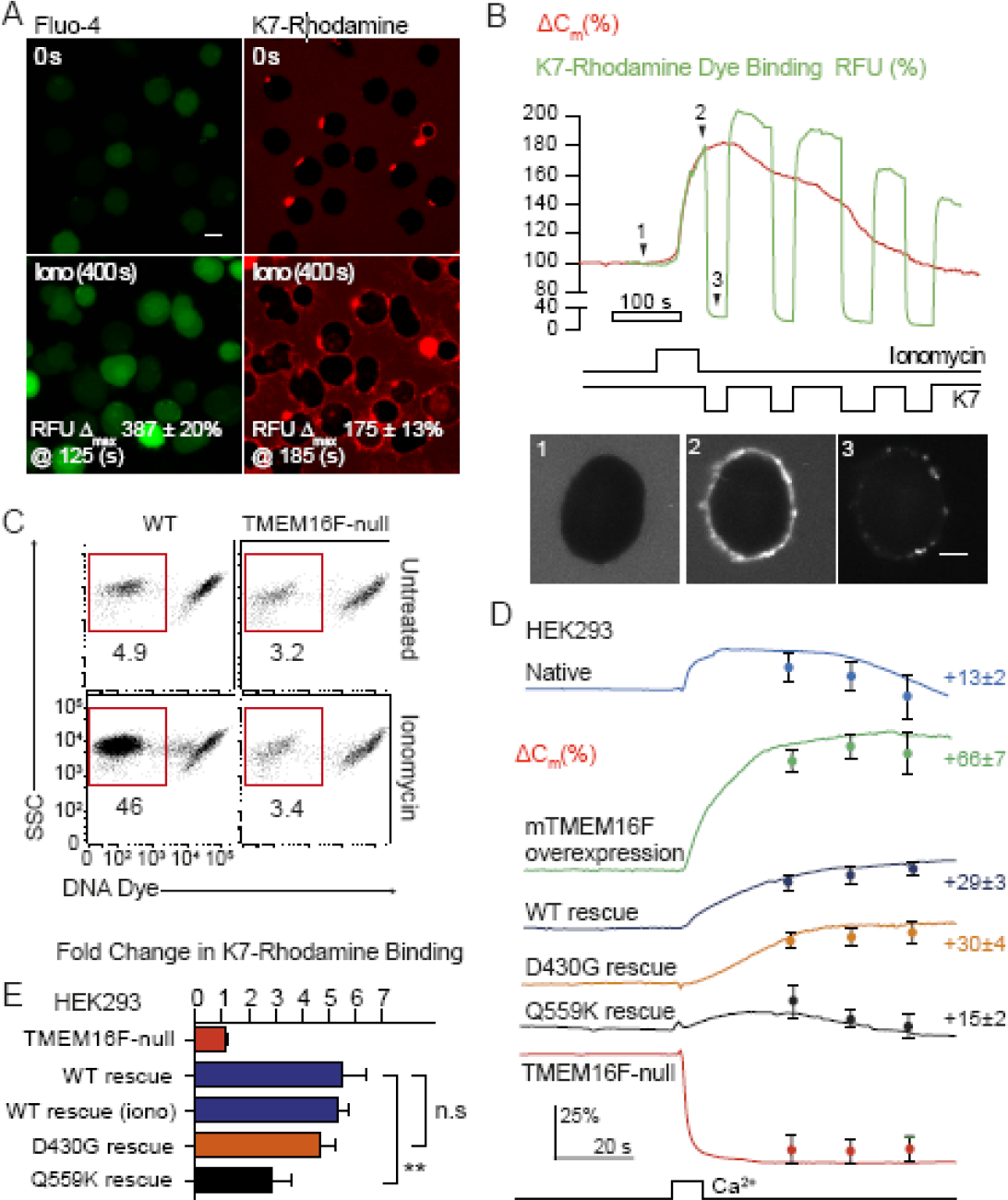
**(A)** Micrograph of Fluo4 and heptalysine-rhodamine (K7r) labelled WT Jurkat cells indicating mean increase and median time to maximal increase for both Ca^2+^ and K7r PS binding. n=4 **(B)** Real-time membrane scrambling and C_m_ were measured with K7r membrane binding with concurrent whole-cell patch-clamp of WT Jurkat T cells using standard solutions following ionomycin treatment. Images are of K7r binding to the cell at indicated time points with scale bar representing 10 urn. Experimental traces from this cell are shown. **(C)** WT Jurkat T cells were treated for 15 min with ionomycin in Ringer and stained with DNA (Viability) eFluor®780 dye (eBioscience). Subcellular sized events were gated and apoptotic bodies distinguished from microvesicles by DNA dye uptake. Numbers are percentage of total events. **(D)** HEK293 TMEM16F-null cells transfected with WT, D409G or Q559K mouse TMEM16F. Cells were imaged before and after Ca^2+^ transients (NCX1 reverse exchange or ionomycin when indicated), and the percentage change in C_m_ was calculated for each group. (**p ≤ 0.01) Single recordings displayed with mean values and SEM at 30, 45 and 60 s after stimulus Mean values with SEM of n>7. **(E)** Fold change in K7r binding for cells described in (D). n>7 for each group

In order to detect the fate of the shed membrane we treated wild-type and TMEM16F-null Jurkat T cells with Ca^2+^ ionophore for 10 minutes, then stained the cultures with Fluor®780 dye which binds to DNA. Flow cytometry was used to analyse subcellular-sized events (**Figure 5C**). All cultures contained similar levels of subcellular, DNA-containing particles. These are likely to be apoptotic bodies, as these cell cultures contain a small fraction of apoptotic cells (data not shown). However, Ca^2+^ ionophore treatment of WT but not TMEM16F-null Jurkat T cells resulted in a large increase of DNA-negative subcellular events, compatible with the release of numerous vesicles.

We next examined to what extent expression of TMEM16F and TMEM16F mutants, reported to have modified scrambling activity (Suzuki et al., 2010; Yang et al.), could rescue the TMEM16F-null phenotype. To do so, we employed the same CRISPR/Cas9 system to create a TMEM16F-null HEK293 cell line that constitutively overexpresses the NCX1 Na/Ca exchanger (TMEM16F-null HEK293-NCX1 cells), which can be used to generate transient Ca^2+^ elevations without ionomycin. As in Jurkat cells, Ca^2+^ influx in TMEM16F-null HEK293 cells caused rapid MEND (**Figure 5D**, red trace), and both membrane expansion (**Figure 5D**) and PS exposure (**Figure 5E**) could be restored by expressing mouse TMEM16F and its mutants. The D430G mutant has been reported to have constitutive PS scrambling activity (Suzuki et al., 2010). However, we found no clear difference between WT-rescue and D430G mutant-rescue in phospholipid scrambling similar to that which has been described in more recent reports (Yang et al., 2012). Additionally we demonstrate no difference in membrane expansion for the D430G mutant versus WT rescue. Interestingly the Q559K mutant, which alters ion selectivity and slows activation kinetics (Yang et al., 2012) had diminished both PM scrambling and membrane expansion responses. However, the residual function of the Q559K mutant was adequate to prevent the MEND response seen in TMEM16F-null cells.

### PD-1 traffics with Ca^2+^-activated membrane movements

To address the specificity of the large-scale trafficking events activated by Ca^2+^, we next examined changes in the surface expression of T cell surface proteins via antibody staining. In WT Jurkat T cells there was a striking reduction in surface expression of the integral membrane protein PD-1 following Ca^2+^-ionophore treatment (**Figure 6A**), compared to a number of other PM proteins. We used a PD-1/GFP fusion protein to demonstrate that this decrease in surface expression was due to loss of the molecule from the cells through membrane shedding, as both PD-1 surface staining and PD-1/GFP fluorescence were reduced upon ionophore treatment (**Figure 6B**). The extent of PD-1/GFP loss was not as great as the loss of PD-1 surface staining (**Figure 6C**), as some of the PD-1/GFP is present intracellularly (data not shown, PD-1/mCherry shown in **Figure 6E**). Shed membrane containing antibody stained PD-1 and PD-1/GFP could be isolated from the supernatant of PD-1/GFP-expressing cells 15 min after ionomycin treatment (**Figure 6D**). Using a PD-1/mCherry fusion protein, we were able to resolve these PD-1 rich vesicles by Structured Illumination Microscopy (SR-SIM) (**Figure 6E**, **Figure 6-figure supplement 1**).

**Figure 6.**
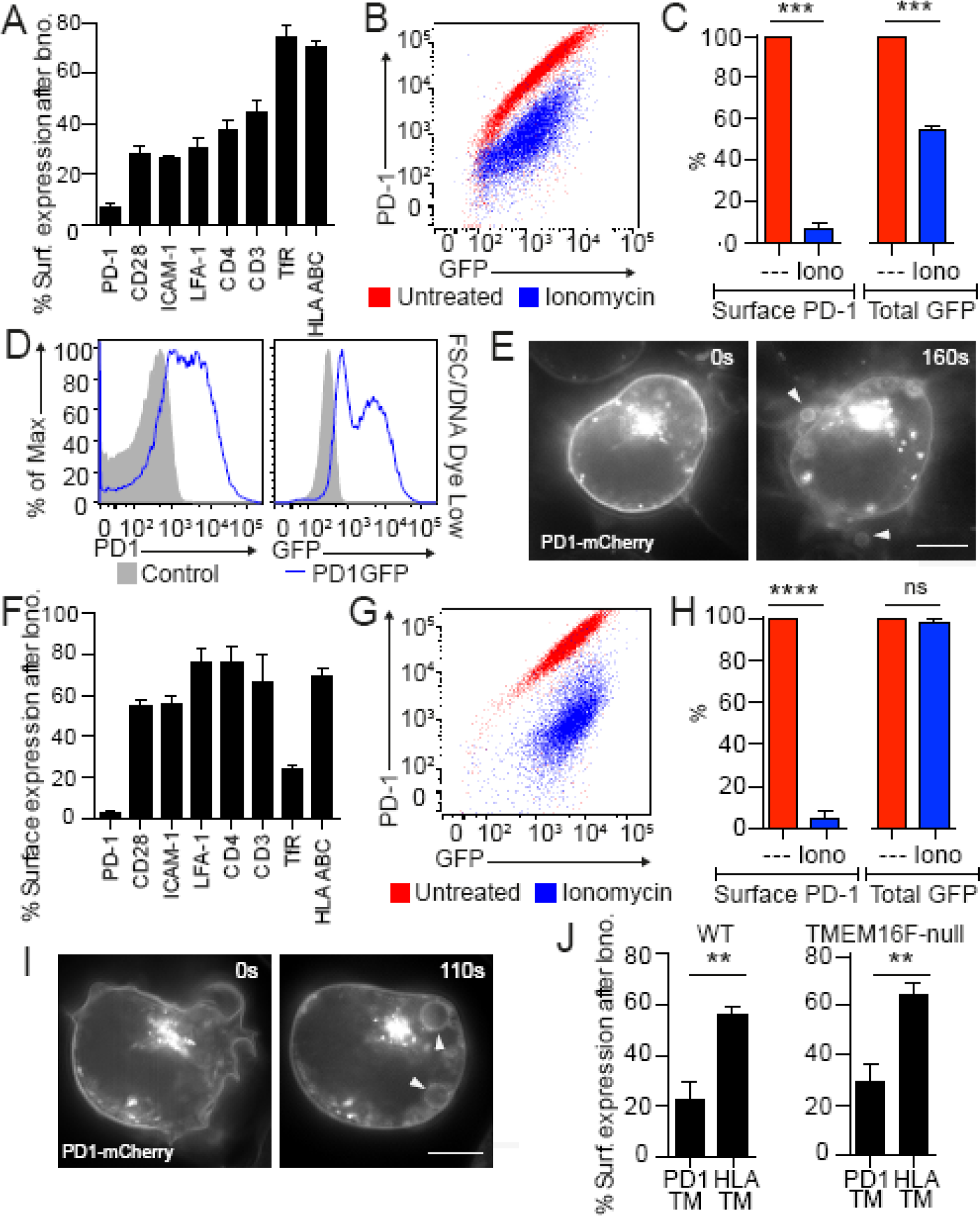
**(A)** WT Jurkat T cells expressing PD-1 were treated with ionomycin for 15 min in Ringer solution and stained for expression of surface molecules. Data are percentage of initial surface expression following ionomycin treatment. **(B)** and **(C)** PD1/GFP chimera was expressed on WT Jurkat T cells. Surface PD-1 and total GFP with and without ionomycin treatment are shown. **(D)** Jurkat WT control and WT PD-1/GFP chimera expressing cells were stimulated with ionomycin. Vesicles were gated for as subcellular sized events with low DNA dye uptake. PD-1 and GFP expression on shed vesicles is shown. **(E)** PD1-mCherry chimera was expressed in WT Jurkat T cells and imaged by SR-SIM microscopy before and after ionomycin treatment. Arrows indicate PD1-bearing microvesicles. **(F)** TMEM16F-null cells were treated and stained as in **(A)**. **(G)** and **(H)**, PD1GFP chimera was expressed in TMEM16F-null Jurkat T cells. Surface PD-1 and total GFP with and without ionomycin treatment are shown. **(I)**, PD1-mCherry chimera was expressed in TMEM16F-null Jurkat T cells and imaged by SR-SIM microscopy before and after ionomycin treatment. **(J)**, PD1HLA-A2 chimeras were produced and expressed in WT and TMEM16F-null Jurkat T cells. Percentage surface expression remaining after treatment with ionomycin is shown. (**p ≤ 0.01, ***p ≤ 0.001 ****p ≤ 0.0001). (For E and I, scale bar is 5 urn; For A, C, F, H, K For A, C, F, H, K, FACS data is for three independent repeats employing >10,000 cells per condition. Data points give mean values with SEM.

When we examined PD-1 surface expression in TMEM16F-null cells, detected by antibody staining, we found this also substantially decreased following Ca^2+^-ionophore treatment (**Figure 6F**). In this case, however, the PD-1/GFP chimera was not lost from the cells when we quantitated GFP fluorescence after Ca^2+^-stimulation (**Figure 6G-H**). Indeed, we could detect the PD-1/mCherry chimera in large intracellular vesicles (**Figure 6I**, **Figure 6-figure supplement 2**) demonstrating that it had been lost from the PM inwardly through the process of MEND. In order to examine which part of the PD-1 molecule controlled its trafficking, we constructed chimeras between PD-1 and HLA-A2, as HLA staining did not change in response to an increase in intracellular Ca^2+^ (**Figure 6A, F**). **Figure 6J** shows that the transmembrane domain of PD-1 targeted its sorting into shed membrane and MEND endosomes, in WT and TMEM16F null cells respectively.

## Discussion

Reconstitution experiments using a recombinant fungal TMEM16 in lipid vesicles demonstrate that this protein can function both as an ion channel and as a phospholipid scramblase that facilitates movement of phospholipids between the inner and outer PM monolayers (Malvezzi et al.). Our results in intact cells show that TMEM16F also functions as a cytoplasmic Ca^2+^ sensor for large-scale PM expansion that is followed by extensive PM shedding. Deletion of TMEM16F blocks both scrambling and shedding and shifts this Ca^2+^ response from PM expansion to MEND. These results provide for the first time a mechanism by which lipid scrambling may account for the diverse phenotypes associated with changes of the expression and function of this channel.

### How are TMEM16F-induced membrane expansion, ion conduction and PS exposure related?

PM expansion occurs within nearly the same time course as TMEM16F conductance changes (**Figure 2A**). Both PM expansion and TMEM16F conductance are blocked by cytoplasmic polyamines (**Figure 3**), physiological metabolites long known to have diverse signalling functions that can include block of anionic phospholipid scrambling (Bucki et al., 1998). Importantly, PM scrambling also occurs with similar kinetics to PM expansion (**Figure 5B**), and both PM expansion and scrambling are reduced by a mutation (Q559K) that affects the selectivity and activation kinetics of TMEM16F (**Figure 5D**). Together, these results suggest that the Ca^2+^-dependent conformational changes that activate TMEM16F conductance also trigger PM expansion. One possibility therefore is that phospholipid scrambling by TMEM16F is the trigger for PM expansion. The appearance of specific phospholipids in the outer monolayer or the loss of specific lipids from the inner monolayer might be the immediate triggers. An alternative is that TMEM16F conformational changes activate large-scale exocytosis and that PS exposure reflects fusion of vesicles containing randomized phospholipids (Mirnikjoo et al., 2009; Lee et al., 2013). In an accompanying article we address these questions, as well as whether PM expansion in these experiments indeed reflects bona fide exocytic events.

### How does TMEM16F prevent endocytosis?

From a cell biological perspective, it appears fundamental that TMEM16F activity determines whether large amounts of membrane are added to or removed from the PM in response to Ca^2+^ influx. Specifically, the occurrence of PM expansion blocks the occurrence of endocytosis. This may be a consequence of changing phospholipid distributions in the PM or possibly also a consequence of rapid changes of the protein content of the PM. The TMEM16F mutant Q559K, which has altered ion selectivity and slow activation kinetics (Yang et al., 2012), permits only a minimal amount of membrane expansion in response to Ca^2+^ influx, but it almost completely blocks endocytic responses. In other words, a minimally active TMEM16F is sufficient to block endocytosis. An explanation suggested by previous work (Fine et al., 2011) is that MEND relies on membrane ordering into domains. TMEM16F activity presumably disrupts ordering of the outer PM monolayer and thereby disrupts MEND. Alternatively, the addition of new membrane might dissipate lateral membrane tension that in some way supports MEND. A third possibility is that signalling mechanisms mediated by TMEM16F conformational changes control both large-scale membrane insertion and loss of PM via mechanisms other than canonical SNARE mediated fusion and clathrin mediated endocytosis. A mechanism for extensive PM expansion without involvement of SNAREs is suggested in an accompanying article (Fine, XXXX).

### Role of membrane traffic in membrane repair

It has long been recognised that cells can rapidly repair mechanical damage to their PM (Wilson, 1924). When the PM is damaged, for example by a physical lesion or a pore-forming toxin, the cytoplasmic Ca^2+^ concentration can rise by more than a hundred-fold, leading to Ca^2+^-dependent recruitment of endomembrane and membrane repair (Bi et al., 1995; Cooper and McNeil, 2015). It seems probable that these same processes will become activated when an ionophore induces a large elevation of cytoplasmic Ca^2+^. Similar to our present experiments using TIRF microscopy to visualize fusion events, the repair process has been analysed extensively in fibroblasts after long-term labelling of the secretory pathway with fluorescent dyes (Togo et al., 1999; Togo et al., 2003). While those studies clearly support the occurrence of classical SNARE-mediated membrane fusion events, the extent of dye destaining that occurs locally at a wound amounts to only 4 to 8% of the local membrane labelling (Togo et al., 2003). Membrane damage and its Ca^2+^-dependent repair via membrane trafficking mechanisms have also been extensively studied in muscle (Cooper and McNeil, 2015). Mutations in both dysferlin (Bansal et al., 2003) and TMEM16E (Griffin et al., 2016), which is closely related to TMEM16F and highly expressed in muscle, cause muscular dystrophies with limitations in membrane repair. Thus, membrane repair may clearly be one physiological function of Ca^2+^-stimulated PM expansion, dependent on TMEM16F. Recently, endocytosis as well as exocytosis has been suggested to play important roles in membrane repair (Idone et al., 2008; Jimenez et al., 2014). Expression or activity of TMEM16F might therefore control the balance between these processes. The observation that the PD-1 transmembrane region associates with membrane that is either shed or endocytosed, depending on TMEM16F activity, suggests that subsets of the PM, i.e. membrane domains, take part in Ca^2+^ stimulated trafficking in a specific manner.

### Role of shed vesicles

The shedding of PM as ectosomes, following TMEM16F activation, may also have multiple physiological and pathological functions (Minciacchi et al., 2015; Zmigrodzka et al., 2016) (**Figure 5, 6**, **Figure 5-figure supplement 1**). PS-expressing vesicles have been proposed to act as a catalyst for bone mineralization (Bevers and Williamson, 2016), and Scott syndrome (TMEM16F-null) cells do not produce these, because they do not expose PS (Bevers et al., 1992). This can explain why TMEM16F-null mice have defective embryonic bone formation (Ehlen et al., 2013). In T cells, we have detected vesicles shed from T cells containing the immune inhibitor PD-1, so vesicle shedding could be one mechanism by which PD-1 is down-regulated, preventing T cell exhaustion. Hu and colleagues (Hu et al., 2016) detected TMEM16F in endosomes that are recruited to the immunological synapse in T cells and postulated that PD-1 down-regulation was due to PD-1 endocytosis leading to degradation. However, our data suggest that recruitment of vesicles to the immune synapse followed by shedding of PD-1 is an intriguing and therapeutically important alternative possibility.

Vesicle shedding from T cells can be a localized phenomenon, for example following TCR triggering (Kumari et al., 2014). Our study is limited in that we have used a T cell line which is amenable to patch-clamp experiments, and a non-physiological Ca^2+^ ionophore stimulus to induce whole cell responses so that we can measure bulk PM changes via capacitance changes. Future studies of the role of TMEM16F measuring local vesicle shedding in primary T cells following receptor triggering are therefore warranted.

## Acknowledgments

HLA-A2 cDNA was a gift from Dr. Martin Pule, UCL. lentiCRISPR v2 plasmid was a gift from Feng Zhang, MIT. We thank David Escors, Djamil Damry, Mehdi Barachian, Mei-Jung Lin and Christopher Tie for help with this project and Dr. Criss Hartzell (Emory) for WT mouse TMEM16F plasmid. CB was supported by The Rosetrees Trust (M155), MRC Doctoral Training Grant (1132770), and the UCL Bogue Fellowship. DH and MF were supported by National Institutes of Health, USA (HL119843, T32DK007257). MFRH and PMP were supported by Medical Research Council (MR/K015826/1) and Biotechnology and Biological Sciences Research Council (BB/M022374/1).

The Authors declare no competing financial interests.

CB designed and performed experiments, wrote manuscript

MF designed and performed experiments, wrote manuscript

PMP designed and performed experiments

JS sequenced genomicTMEM16F alleles

MT designed and isolated PD-1 constructs

RH designed experiments

MKC designed experiments, wrote manuscript

DH designed and performed experiments, wrote manuscript

## Materials and Methods

Data analysis was performed in Prism (GraphPad) or SigmaPlot (Systat) unless otherwise stated.

### Cell lines, solutions and reagents

Jurkat E6.1 T cells (ECACC) were grown in RPMI-1640 (Sigma-Aldrich) while HEK293 cells (Invitrogen) were grown in DMEM (Sigma-Aldrich). All media were supplemented with 10% FCS (Invitrogen), 2mM L-glutamine (Invitrogen) with 100 U/mL penicillin and 100 μg/mL streptomycin (Invitrogen). Standard cytoplasmic solution contained (in mM) 145 KCl, 10 HEPES, 0.5 EGTA, 0.25 CaCb, 8 MgATP, 2 TrisATP and 0.2 GTP, adjusted to pH 7.4. Standard extracellular solution contained (in mM) 145 NaCl, 10 HEPES, 2 CaCl_2_ and 2 MgCl_2_, adjusted to pH 7. K^+^-free *N*-Methyl-D-glucamine (NMG) aspartatic acid (Asp) solution (NmAs) contained (in mM) 145 NMG, 10 HEPES, and 0.5 EGTA, adjusted to pH 7 with Asp; 2 CaCl2 or 2 MgCl2 were added when solution was used on the extracellular side while 0.25 and 0.5 were used, respectively, on the cytoplasmic side. Sucrose solution contained (in mM) 240 sucrose, 5 HEPES, 0.3 EGTA, and 5 KCl at pH 7; 2 CaCl_2_ or 2 MgCl_2_ were added when solution was used on the extracellular side while 0.25 and 0.5 were used, respectively, on the cytoplasmic side. 5μM ionomycin free acid (Calbiochem) was used unless otherwise stated. Other reagents used were: Lipofectamine 3000 (Life Technologies) for transient transfection protocols; STIM1-YFP (a generous gift Jen Liou, UTSW); 5μM FM 4-64 (Life Technologies), 400nM tetanus toxin L-chain (Alpha Universe LLC), 3μM latrunculin A (Sigma-Aldrich), 10μM phalloidin (Tocris), 3μM myristoylated dynamin inhibitory peptide (Tocris), 100nM LysoTracker Red DND-99 (Life Technologies), 50-500μM spermine (Sigma-Aldrich), and 100μg/ml (0.01%) Trypan Blue (Sigma-Aldrich). Rhodamine-conjugated hepta-lysine (rhodamine-KKKKKKK-amide; K7r) was prepared by Multiple Peptide Systems (NeoMPS, Inc.) and used at 3μM. Compared to Annexin V, K7r was advantageous for real-time imaging as membrane binding is significantly faster, and K7r binding does not require Ca^2+^.

### Whole-cell patch clamp with confocal imaging

Patch clamp recording of cell electrical parameters was performed as described previously (Yaradanakul et al., 2008) using Capmeter v6.3 (Wang and Hilgemann, 2008) with National Instruments digital acquisition boards and Axopatch 200B or 1D patch clamp amplifiers. Square-wave voltage perturbation (20 mV; 0.5 kHz) was employed, input resistances were 2-9 MΩ, and the apparent cell resistances were 0.1-2 GΩ. External solutions were temperated to 35-37°C in parallel gravity-fed solution lines with outlet flow velocities of 2 to 5 mm/s. When confocal imaging and electrophysiology were combined, a Nikon TE2000-U microscope; 60× oil immersion, 1.45-NA objective paired with the EZ-C1confocal system (Nikon) containing a 40-mW 163-Argon laser (Spectra Physics; Newport Corporation) operating at 488 nm at 7.5% maximum capacity for FM4-64 and YFP recordings and a 1.5mW Melles Griot cylindrical HeNe laser at 543nm at 55% of maximum capacity for K7r, LysoTracker Red and Trypan Blue. Emission filters were set to either 500-540nm or 580LP. Typical resolution was 512 × 512, set with a pixel dwell time yielding <1-s exposure times per frame and a pinhole of 150μm. Image analysis was performed using either the EZ-C1 v3.9 (Nikon Instruments) or ImageJ (NIH). When imaging multiple fluorophores, sequential imaging was used to minimize spectral overlap. Photobleaching was negligible for these experiments. For independent confocal imaging of intact cells, 800μl of media containing Jurkat cells was transferred to a glass bottom 4-chamber dish. Warmed ionomycin solution was rapidly added at a 5:1 concentration and cells were imaged as described. For patch clamp, intracellular calcium stimulus was triggered by direct perfusion of the Ca^2+^ ionophore ionomycin for Jurkat and HEK293 cell lines. HEK293 cells stably expressing the cardiac Na/Ca exchanger (NCX1.1) were stimulated via reverse Na/Ca exchange activated by the presence of 40mM Na^+^ on the cytoplasmic side and application of 2mM Ca^2+^ on the extracellular side for 5 to 20 s (Yaradanakul et al., 2008). Functional responses of HEK293 cells to ionomycin application or reverse Na/Ca exchange activation for 5 to 20 s were not significantly different (**Figure 5E**).

### CRISPR-Cas9 knockdown of TMEM16F

The lentiCRISPR v2 plasmid was used as described (Sanjana et al., 2014). The guideRNA sequence targeting TMEM16F Exon 2 was 5’-TCAGCATGATTTTCGAACCC-3’ and TMEM16F Exon 5 was 5’-TGTAAAAGTACACGCACCAT-3’. HEK293 cells were transiently transfected with plasmid using Lipofectamine (Life Technologies) and the lentivirus was collected from the supernatant. Jurkat T cells and HEK293 cells were transduced with lentivirus produced using the gag-pol expression vector p8.91 and VSV-G envelope vector pMDG as described previously (Zufferey et al., 1997). 48 hours after transduction cells were selected in 0.5μg/ml Puromycin for Jurkat and 5μg/ml Puromycin for HEK293 before single-cell clones were isolated. Cells stably expressing Cas-9 and gRNA after transduction were frozen after initial expansion and aliquots thawed as needed to avoid accrued off-target effects resulting from long-term culture. Gene editing was confirmed by sequencing (Beckman Coulter) (**Figure 1-figure supplement 1**). Two different primer sets were used to amplify Exon 2 (Set 1 and 2) and one for Exon 5 (Set 3) and the amplicons sequenced using the same primers. Primer sequences were as follows (all 5’ - 3’):

Set 1 FW GTGCAGGTTCATGCTTCATTT,
Set 1 RS GCACAGTCCCTGCAATAAGA,
Set 2 FW CCCGGTGCTGCTGATTTA,
Set 2 RS GGATCTACAGCCATTGAAGGAA,
Set 3 FW AGTGGTGGTCTCTGTATTGTTT,
Set 3 RS AAACAGCAGGTTCCAAATTACC.

#### Proteomic analysis of Jurkat WT and CRISPR-Cas9 knock-out cells

Cells were subjected to Dounce homogenation on ice in 250 mM sucrose/10 mM Tris pH 7.4 with cOmplete protease inhibitor (Roche). Cell debris was pelleted and the supernatant was spun at 2000g for 60 min to pellet membrane. Membranes were washed and reconstituted with 4 X Laemmli buffer for SDS-PAGE. Protein bands were excised and digested overnight with trypsin (Pierce) following reduction and alkylation with DTT and iodoacetamide (Sigma-Aldrich). The samples underwent solid-phase extraction cleanup with Oasis HLB plates (Waters) and the resulting samples were analyzed by LC/MS/MS, using an Orbitrap Fusion Lumos mass spectrometer (Thermo Electron) coupled to an Ultimate 3000 RSLC-Nano liquid chromatography systems (Dionex). Raw MS data files were converted to a peak list format and analyzed using the central proteomics facilities pipeline (CPFP), version 2.0.3 (Trudgian et al., 2010; Trudgian and Mirzaei, 2012). Peptide identification was performed using the X!Tandem (Craig and Beavis, 2004) and open MS search algorithm (OMSSA) (Geer et al., 2004) search engines against the human protein database from Uniprot, with common contaminants and reversed decoy sequences appended (Elias and Gygi, 2007). Label-free quantitation of proteins across samples was performed using SINQ normalized spectral index Software (Trudgian et al., 2011).

### Total Internal Reflection Fluorescence Microscopy (TIRFM)

Methods were adapted from (Zhang et al., 2007; Li et al., 2008). Jurkat T lymphocytes were incubated with 5μM FM4-64 in RPMI for 3 hours. Cells were then washed thoroughly and rested for 20 min in Ringer solution before imaging in 8 well glass bottom μ-slides (1.5H, ibidi). Ionomycin was added to wells at a final concentration of 5μM, 5s before recording began. Imaging was performed in an ElyraPS.1 inverted microscope (Zeiss), equipped with an EMCCD camera (iXon 897, Andor), in TIRF mode. Acquisition was done using a 100x TIRF objective (Plan-APOCHROMAT 100x/1.46 Oil, Zeiss), using the definite focus (Zeiss) to avoid drift in Z. The frame rate was 10fps. Putative fusion events were defined as complete puncta destaining to background level, with duration of destaining ≤1.5 s (15 frames), with no sideways movement of puncta and were manually counted, while puncta number was quantified using QuickPALM (Henriques et al., 2010) and verified using ImageJ particle analysis after implementing Huang’s fuzzy thresholding method using Shannon’s entropy function and watershed algorithm (Huang, 1995).

### Chimeric proteins, lentiviral vector constructs and production

All chimeric proteins were produced using overlap-extension PCR (Ho et al., 1989) and cloned into a single (pSIN) or dual (pDUAL) promoter vector. In fluorescent constructs a ‘GS’ linker separated PD-1 and fluorophore. Primers used to make PD1/GFP were PD1-FW, PD1GFP Sense, and PD1GFP Antisense GFP-RS. PD-1 mCherry was made in the same way using the primers PD1FW, PD1mCherry Sense, PD1mCherry Antisense and mCherry RS. PD1HLA Chimers were made with PD1-FW and HLA-A2-RS. For PD-1HLA-A2 chimeras, the extracellular portion was always PD-1 and the intracellular HLA-A2. The TM regions were the only sequences to vary between the two constructs. The transmembrane regions were predicted using the TMPred algorithm (https://embnet.vital-it.ch/software/TMPRED_form.html. The PD-1 TM was predicted as L168 to L191 and HLA-A2 TM I410 to W435. The chimera with PD-1 transmembrane was made with PD1TM Sense and PD1TM Antisense primers. The chimera with HLA-A2 transmembrane was made with HLATM Sense and HLATM Antisense primer. Lentivirus was produced using the gag-pol expression vector p8.91 and VSV-G envelope vector pMDG as described previously. The primer sequences were as follows:

PD1-FW GGGGGGATCCGCCACCATGCAGATCCCACAGGCGCC,
PD1GFP Sense GGCTCCGGCTCCGGCTCCGTGAGCAAGGGCGAGGAGCT,
PD1GFP Antisense GGAGCCGGAGCCGGAGCCGAGGGGCCAAGAGCAGTGTC,
GFP-RS GCGGCCGCTTTACTTGTACAGCTCGTCCATGCCGAGAGTGATCCC,
PD1mCherry Sense GGCTCCGGCTCCGGCTCCGTGAGCAAGGGCGAGGAGGA,
PD1mCherry Antisense GACACTGCTCTTGGCCCCTCGGCTCCGGCTCCGGCTCC,
mCherry RS GGCTCCGGCTCCGGCTCCGTGAGCAAGGGCGAGGAGGA,
HLA-A2-RS CCCCGCGGCCGCTCACACTTTACAAGCTGTGAGAGACACATCAG,
PD1TM Sense TCTGGGTCCTGGCCGTCATCAGGAGGAAGAGCTCAGATAG,
PD1TM Antisense CTATCTGAGCTCTTCCTCCTGATGACGGCCAGGACCCAGA,
HLA TM Sense CAGCCGGCCAGTTCCAAACCATCGTGGGCATCATTGCTGG,
HLA TM Antisense CCAGCAATGATGCCCACGATGGTTTGGAACTGGCCGGCTG.

### Super-resolution structured illumination microscopy (SR-SIM) and Zeiss Airyscan microscopy

SR-SIM and Airyscan imaging was performed at 37°C using Plan-Apochromat 63x/1.4 oil DIC M27 objective in an Elyra PS.1 microscope (Zeiss). For SR-SIM, images were acquired using 5 phase shifts and 3 grid rotations, with 34μm grating period for the 561nm laser and filter set 4 (1851-248, Zeiss). Airyscan z-stacks were obtained at 0.75um steps for a total depth of 12μm. Stacks were obtained every 30sec using a fast piezo controller and Definite Focus 2.0 (Zeiss) to avoid z-drift between time points. Processed images were projected into a single 2-D image based on maximum intensity using ImageJ. SR-SIM images were acquired using an sCMOS camera and highresolution Airyscan images were acquired using the LS880 GaAsP detector, both processed using ZEN software (Zeiss).

### Flow cytometry, antibodies and dyes

Cells were treated with ionomycin for 15 min at 37°C in Ringer solution containing 2 mM CaCl2. Cells were stained for 30 min on ice and flow cytometry was performed using a BD Fortessa (BD Bioscience). Intracellular Ca^2+^ concentration was monitored with 1μM Fluo-4 AM (Life Technologies) loaded in cells for 30 min at 37°C prior to the experiment. Surface exposure of phosphatidylserine was detected using 9ng/ml Annexin V staining in Annexin binding buffer (BioLegend). Microvesicles were distinguished from apoptotic bodies using the DNA (Viability) eFluor^®^ 780 dye (eBioscience). Surface staining for receptors was carried out for 30min on ice using the following antibodies: PD-1 APC (MIH4), ICAM-1 PE (HA58), Transferrin receptor (TfR) PE (M-A712) from BD Biosciences and CD28 PE (CD28.2), LFA-1 PE (HI111), CD4 PE (OKT4), CD3 PE/Cy7 (UCHT1), HLA ABC PE (W6/32) from BioLegend. Data were analyzed using FlowJo Software (Treestar).

---

**Figure 1 - figure supplement 1.**
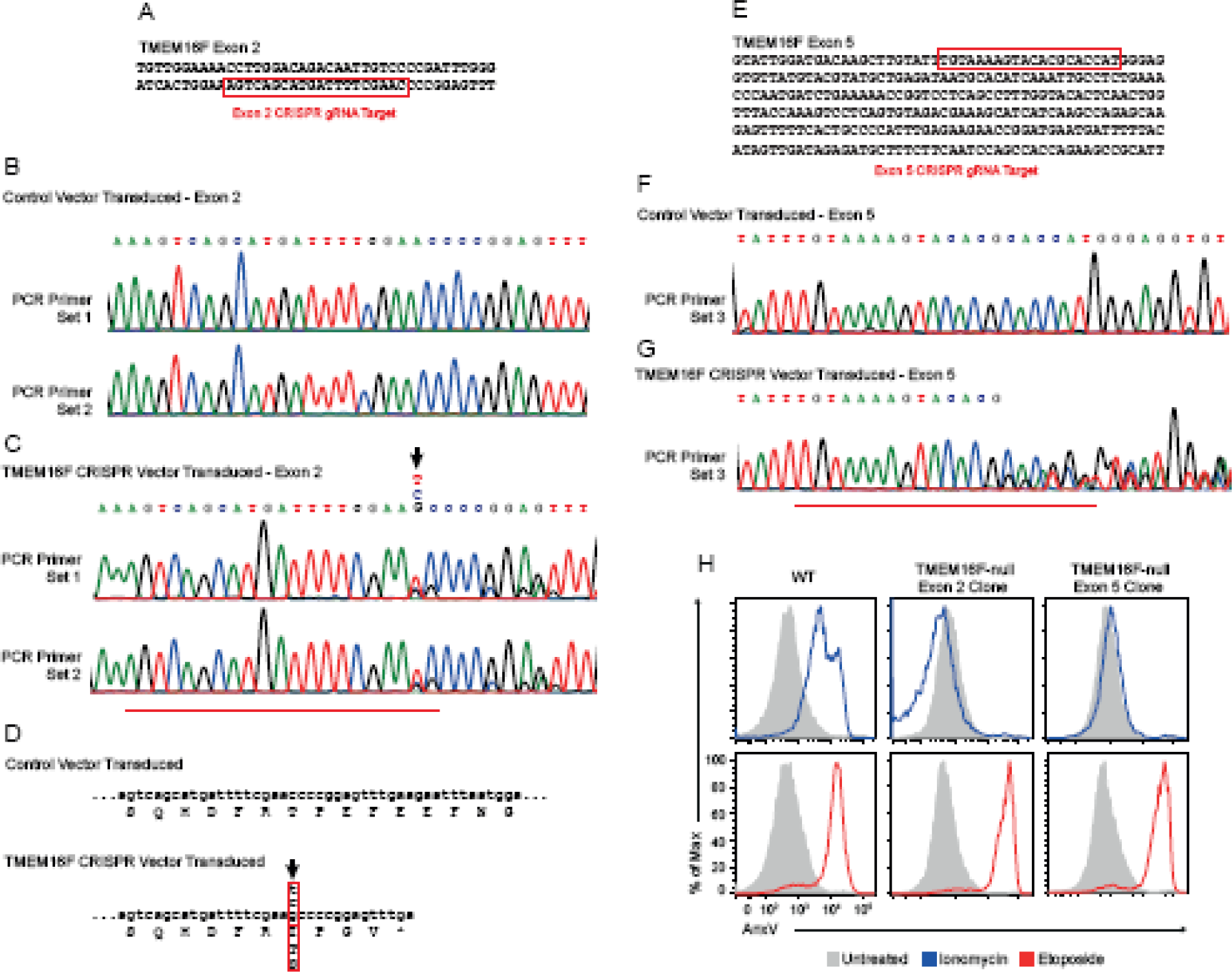
(A) TMEM16F was edited using the lentiviral vector-based CRISPR-Cas9 system. Cells were verified genomically using sequencing analysis and functionally assayed for PS exposure with Annexin V as described in **Figure 4A**. TMEM16F Exon 2 and target sequence is shown. (B) Two sets of primers were used to amplify TMEM16F Exon 2 from vector-only transduced Jurkat T cell genomic DNA. The two amplicons were then sequenced. (C) The same amplicons were sequenced from a clone transduced with TMEM16F Exon 2-targeted CRISPR-Cas9 lentiviral vector. (D) The result of the single nucleotide insertion on the amino acid sequence. A stop codon results 10 nucleotides downstream of the insertion that truncates TMEM16F from 910 to 50 amino acids. (E) TMEM16F Exon 5 and target sequence. (F) One primer set was used to amplify TMEM16F Exon 5 from vector-only transduced Jurkat T cell genomic DNA and sequenced (G) The same amplicon was sequenced from a clone transduced with TMEM16F Exon 5-targeted CRISPR-Cas9 lentiviral vector. The indeterminate sequence 3’ of the target sequence is suggestive of differently mutated alleles and was consistent across repeats. (H) WT, TMEM16F-null cells CRISPR-targeted to either Exon **2** or Exon 5 were treated with 5 μM ionomycin for 15 min or 2 μM Etoposide overnight and stained with Annexin V and compared to untreated cells. WT cells treated with ionomycin show a clear increase in Annexin V binding to exposed phosphatidylserine. Both Exon2 and Exon 5 TMEM16F-null cell lines fail to show Annexin V binding responses during ionomycin treatments, demonstrating that ionomycin-induced scrambling is consistently blocked in cell lines where TMEM16F was targeted for CRISPR-Cas9 excision. Loss of TMEM16F does not interfere with apoptosis-induced scrambling and PS exposure as overnight treatment of Etoposide shows similar increases in Annexin V positive cells for WT and both Exon2 and Exon5 targeted cell lines. Exon2 targeted cell lines were used for further experiments.

**Figure 1 - figure supplement 2 (Table 1).**
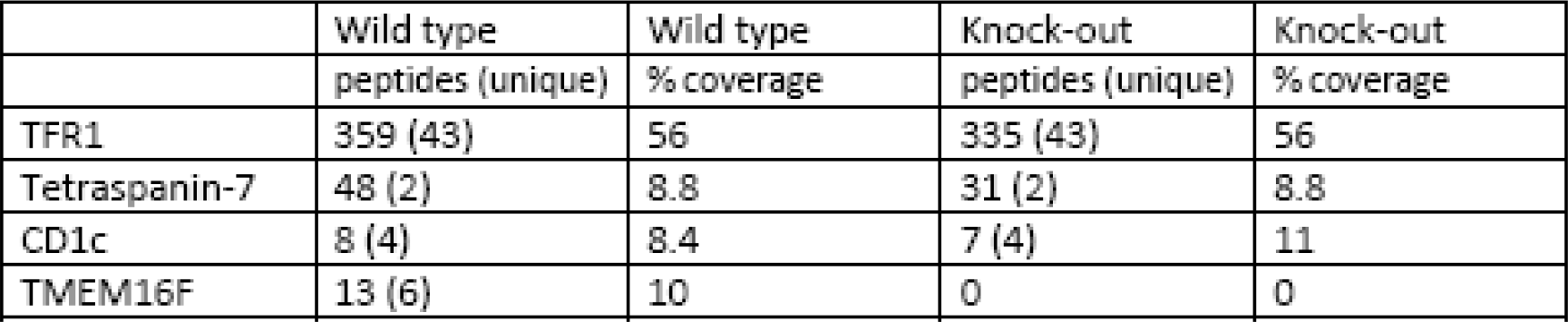
Proteomics analysis of Jurkat WT and CRISPR-Cas9 TMEM16F-null cells. Table describing surface proteins and TMEM16F expression in parental and stably selected Jurkat cell lines.

Figure 2 - figure supplement 1 (video). Ca^2+^-induced exocytosis and shedding probed with FM4-64 in WT Jurkat T cells. WT Jurkat cells were plated in a chambered coverslip in Ringer with 5 μM FM 4-64. 5 μM ionomycin was added at 30 s. The scale bar represents 10μm.

Figure 2 - figure supplement 2. SR-SIM microscopy images of FM4-64 dye distribution before **(A)** and after **(B)** Ca^2+^ influx in TMEM16F-null Jurkat T cells. **(C)** and **(D)** show the time course of MEND in two regions of interest (blue (C) and green (D)). Intracellular vesicles decorated with FM4-64 are visible following MEND, with no evidence of PM permeabilization to FM4-64. The scale bars represent 10 urn.

Figure 5 - figure supplement 1 (video). Ca^2+^ induced PS exposure probed with K7r in WT Jurkat T cells. WT Jurkat cells were plated in a chambered coverslip in Ringer with 3 μM K7r. 5 μM ionomycin was added at 50 s. The scale bar represents 10μm.

Figure 5 - figure supplement 2 (video). Similar to Figure 5 Supplemental 1 Video corresponding the experiment quantified in Figure 5A. Ca^2+^induced PS exposure probed with K7r in WT Jurkat T cells. WT Jurkat cells were pre-incubated with 3μM Fluo-4 (green) and plated in a chambered coverslip in Ringer with 3μM K7r (red). 5μM ionomycin was added at 80 s. The scale bar represents 10 μm.

Figure 6 - figure supplement 1 (video). In WT Jurkat T cells PD-1 is shed following ionomycin treatment. PD1-mCherry expressing WT Jurkat T cells were imaged by SR-SIM Microscopy. Ionomycin was added to the opposite side of the chambered cover slip 10s before the video starts. Scale bar represents 10μm.

Figure 6 - figure supplement 2 (video). In TMEM16F-null Jurkat T cells PD-1 is endocytosed following ionomycin treatment. PD1-mCherry expressing TMEM16F-null Jurkat T cells were imaged by SR-SIM Microscopy. Ionomycin was added to the opposite side of the chambered cover slip 10s before the video starts. Scale bar represents 10μm.

**Supplemental Table 1 (Figure 1).** Supplemental Table 1 (Figure 1)

|  | Wild type | Wild type | Knock-out | Knock-out |
| --- | --- | --- | --- | --- |
|  | peptides (unique) | % coverage | peptides (unique) | % coverage |
| TFR1 | 359 (43) | 56 | 335 (43) | 56 |
| Tetraspanin-7 | 48 (2) | 8.8 | 31 (2) | 8.8 |
| CD1c | 8 (4) | 8.4 | 7 (4) | 11 |
| TMEM16F | 13 (6) | 10 | 0 | 0 |

